# Antimicrobial activity of the BAM complex-targeting peptides against multidrug-resistant gram-negative bacteria

**DOI:** 10.1101/2024.04.15.589587

**Authors:** Eisaku Yoshihara, Hidefumi Kakizoe, Kazuo Umezawa, Kentaro Wakamatsu, Satomi Asai

**Affiliations:** Department of Laboratory Medicine, Tokai University School of Medicine, Isehara, Kanagawa 259-1193, Japan; Department of Emergency and Critical Care, Tokai University School of Medicine, Isehara, Kanagawa, Japan; Department of Respiratory Medicine, National Hospital Organization Omuta National Hospital, Omuta City, Fukuoka, Japan

**Keywords:** BAM complex, targeting peptides, antimicrobial activity, MDR Gram-negative bacteria

## Abstract

Multidrug-resistant (MDR) Gram-negative bacteria represent a notable public health threat, necessitating the urgent development of new antimicrobial agents. The β-barrel assembly machinery (BAM) complex, crucial for the biogenesis of outer membrane proteins in Gram-negative bacteria, emerges as a promising target for drug development. The BAM complex comprises five proteins (BamA–E), and its functionality might be disrupted by peptides targeting the BamA and BamB interface. We synthesized the peptide LTLR, located in the BamB interaction region with BamA in *Pseudomonas aeruginosa*, and assessed its anti-*P. aeruginosa* activity. This peptide demonstrated potential in enhancing antibiotic efficacy against *P. aeruginosa*. Conversely, LRTL, a scrambled version of LTLR, lacked activity, underscoring the importance of specific target site binding for the peptide’s adjuvant effect. Subsequently, we tested the hypothesis that peptide dimerization enhances its effectiveness. A dimeric peptide, composed of two LTLR sequences, was tested and found to possess bactericidal activity against *P. aeruginosa*, an outcome not observed with the monomeric form. Importantly, this dimeric peptide also showed bactericidal activity against MDR *P. aeruginosa* strains. In summary, our findings offer a foundation for developing novel antimicrobial agents against MDR Gram-negative bacteria.

**IMPORTANCE:** This work demonstrates that (1) the β-barrel assembly machinery (BAM) complex-targeting peptides can exhibit antimicrobial activity and that (2) the dimeric forms of these peptides exhibit bactericidal activity against multidrug-resistant (MDR) strains of *P. aeruginosa* and *A. baumannii*.

## INTRODUCTION

A growing number of bacteria are developing resistance to multiple antibiotics currently in use, posing a global public health threat with the emergence of multidrug-resistant (MDR) bacteria (1–4). Gram-negative bacteria, in particular, are of notable concern in healthcare settings. Consequently, there is an urgent need for the development of novel antimicrobial agents against MDR Gram-negative bacteria. Identifying druggable targets distinct from those of conventional antibiotics is crucial for this endeavor.

Gram-negative bacteria possess an outer membrane pivotal for various essential functions, including nutrient import, cell signaling, and adhesion (5). Within this membrane, the BAM complex plays a critical role in outer membrane protein biogenesis, presenting itself as a promising druggable target (6–13). The primary challenge now lies in inhibiting its function. Our hypothesis is that by inhibiting protein-protein interactions within the BAM complex, which comprises five proteins (BamA–E) (14–16), we can prevent the formation of a functional complex, leading to the inhibition of its activity. To test this hypothesis, peptides targeting the interface between BamA and BamB were employed to obstruct protein–protein interactions. Herein, we investigated these peptides for antimicrobial activity against MDR strains of *P. aeruginosa* and *A. baumannii*.

## RESULTS

### The BAM Complex-Targeting Peptide

Four peptides were designed based on amino-acid sequences within the BamB region that are known to interact with BamA.

### Peptide Activity Targeting the BamA–BamB Interface

We synthesized a peptide containing the LTLR amino acid sequence within the PVLTLRETGAP sequence of the BamB regions interacting with BamA in *P. aeruginosa* (17) and assessed its antimicrobial efficacy against *P. aeruginosa*. Notably, this peptide exhibited no discernible impact on bacterial growth. To explore its potential as an antibiotic adjuvant, we incubated *P. aeruginosa* PAO1 cells with the antibiotic polymyxin B (PMB) in the presence or absence of this peptide. As depicted in Fig. 1, while PMB alone at 0.4 mg/ml showed minimal effect on bacterial growth, its bactericidal activity increased proportionally with peptide concentration, suggesting that the LTLR peptide could augment PMB’s effectiveness, with an approximate ED50 value of 1.9 mM.

**Fig. 1.**
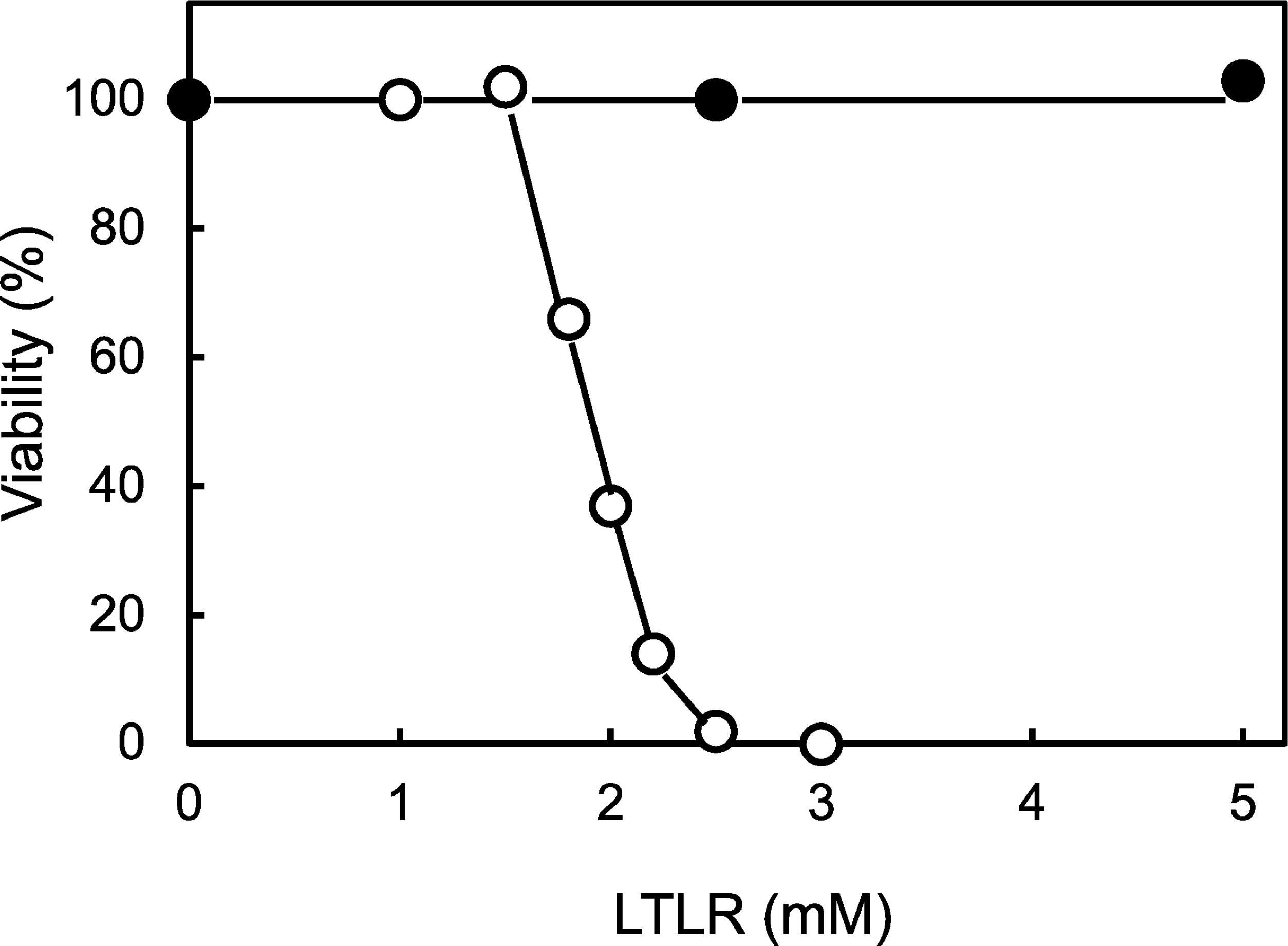
Adjuvant activity of the targeting peptide against *P. aeruginosa*. *P. aeruginosa* PAO1 cells were incubated with polymyxin B (0.4 mg/ml) in the absence or presence of the targeting peptide LTLR (○) or the scrambled peptide LRTL at 1.5 mM (●) and 37 °C for 4 h. Subsequently, cell cultures were diluted with PBS and spread on agar plates. After overnight incubation, the number of colonies was counted. Cell viability was determined using untreated cells as a control.

To ascertain whether the adjuvant effect was specific to the LTLR sequence, we synthesized a scrambled peptide, LRTL, and evaluated its adjuvant potential. Cell viability assays conducted after incubating *P. aeruginosa* PAO1 with PMB (0.4 mg/ml) in the presence or absence of the scrambled peptide revealed no adjuvant activity (Fig. 1), underscoring the importance of the peptide’s specific binding to the target site for adjuvanticity.

Given that membrane permeability influences peptide activity, particularly in traversing the outer membrane to reach its target, we hypothesized that the negatively charged C-terminus of the peptide might hinder permeability due to the outer membrane’s negative charge (5). To test this, we synthesized a peptide with the LTLR amino acid sequence amidated at the C-terminus (designated LTLR’) and evaluated its adjuvant activity. Cell viability assays following incubation of *P. aeruginosa* PAO1 with PMB (0.4 mg/ml) in the presence of varying concentrations of LTLR′ revealed adjuvant activity, with an approximate ED50 value of 0.6 mM, lower than that of the LTLR peptide (Fig. 2). This finding suggests that LTLR′ exhibits enhanced activity compared to LTLR, with the negative charge at the C-terminus adversely affecting peptide permeability through the outer membrane.

**Fig. 2.**
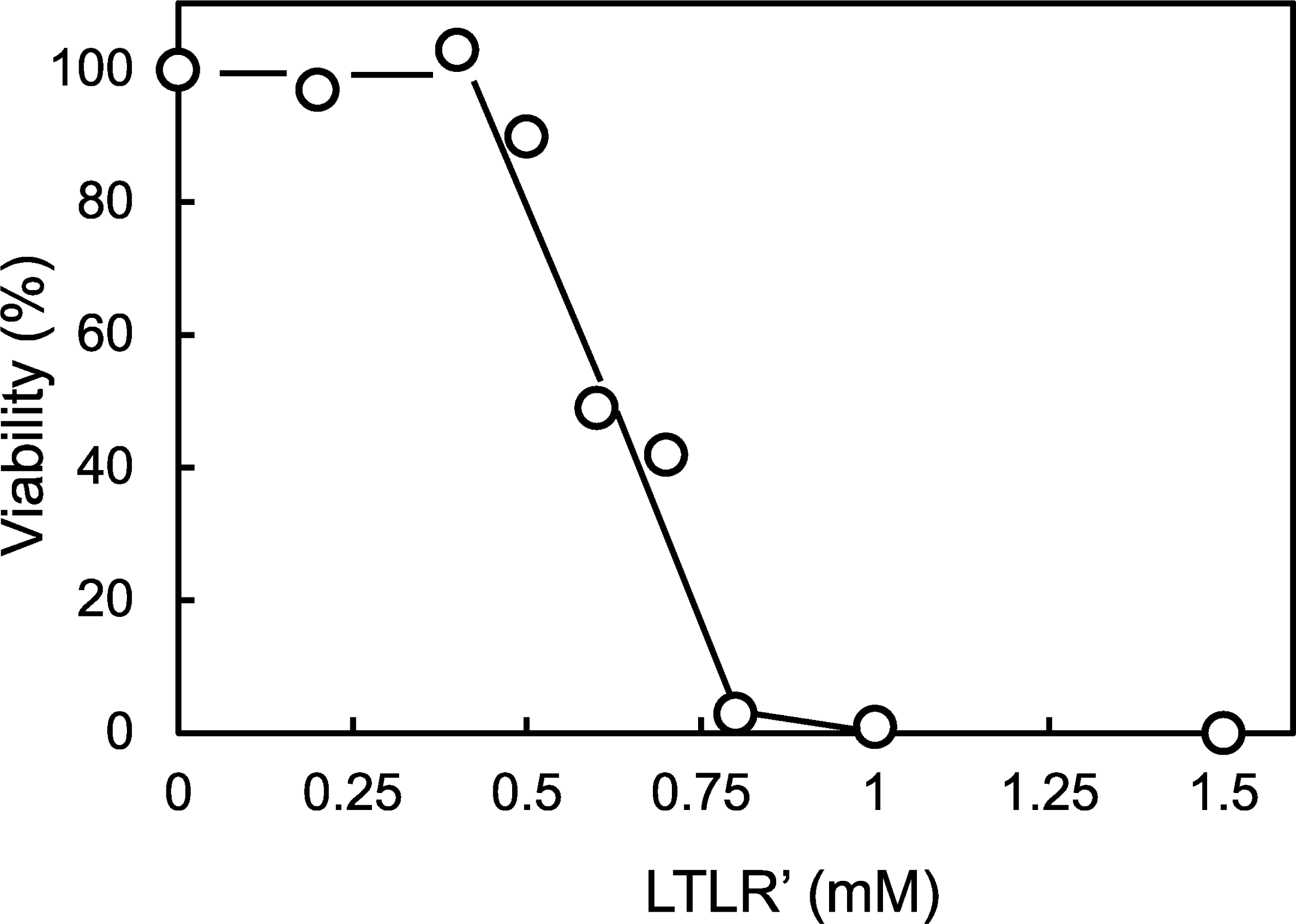
Adjuvant activity of LTLR′ against *P. aeruginosa*. Wild-type *P. aeruginosa* PAO1 were incubated with LTLR′ at different concentrations in the presence of polymyxin B (0.4 mg/ml) at 37 °C for 4 h. Cell viability was determined using untreated cells as controls.

Although these peptides showed adjuvant activity with PMB, they also enhanced the efficacy of other antibiotics. To investigate this further, we employed ofloxacin (OFLX) owing to its distinct mechanism of action compared to PMB and utilized OFLX-resistant *P. aeruginosa* nalB strains. Here, we used *P. aeruginosa* nalB, which highly expresses the MexAB-OprM efflux pump, to examine whether this efflux pump may affect the peptide activity. After incubating *P. aeruginosa* nalB with different OFLX concentrations in the presence or absence of 5 mM LTLR′, cell viability assays were conducted. The results (Fig. 3) revealed that LTLR′ lowered the ED50 values of OFLX from approximately 1.2 mg/ml to approximately 0.1 mg/ml, indicating its potential to enhance OFLX activity. This observation suggests that LTLR′ is not a substrate extruded by the MexAB-OprM efflux pump.

**Fig. 3.**
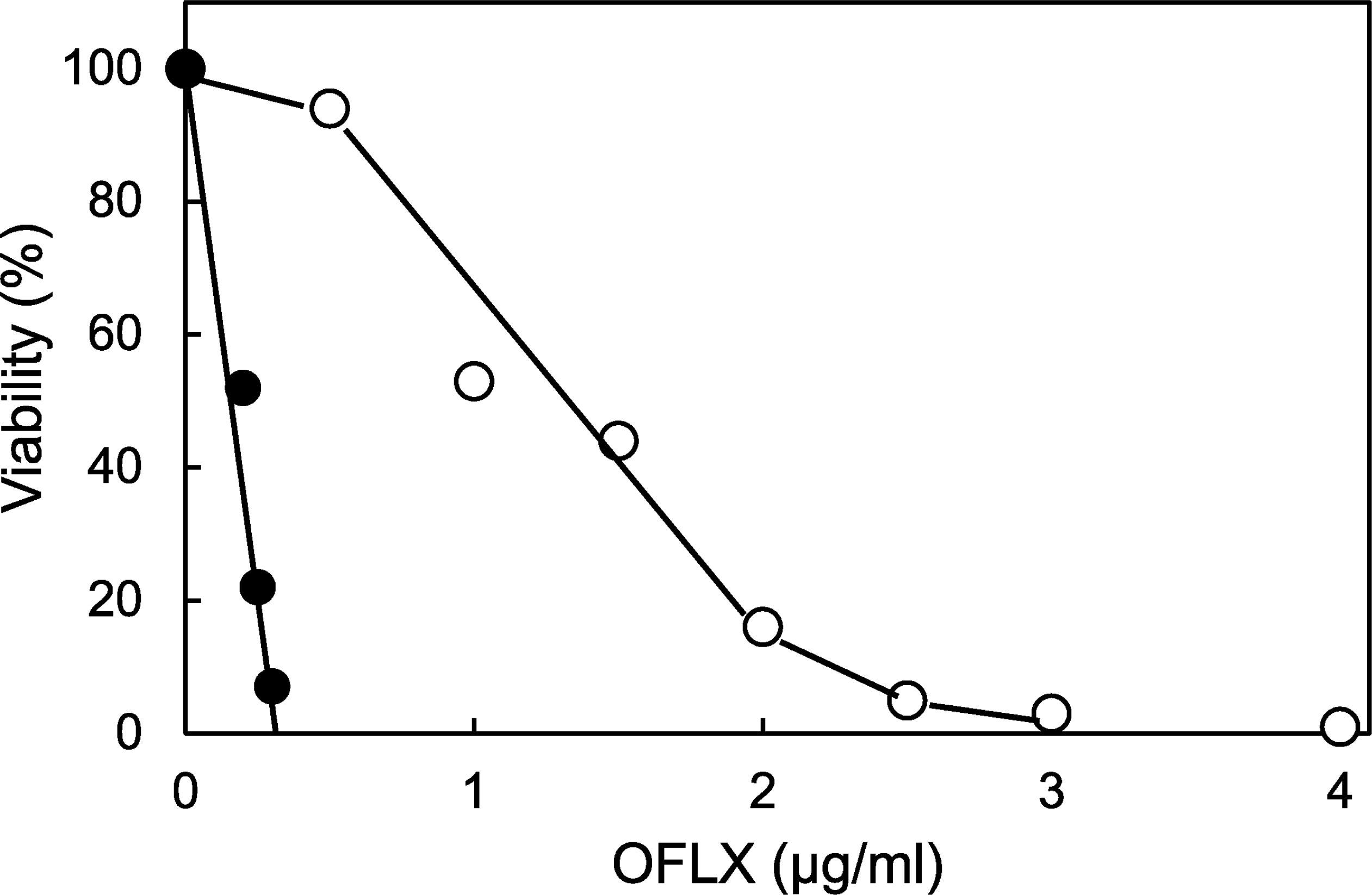
Adjuvant activity of LTLR′ against *P. aeruginosa* nalB strain. *P. aeruginosa* nalB strains were incubated with OFLX at different concentrations in the absence (○) or presence (●) of 5 mM LTLR′ at 37 °C for 4 h. Cell viability was determined using untreated cells as a control.

### Activity of the Dimeric Peptide

We further explored the potential enhancement of peptide activity through peptide modification. Considering that dimerization might amplify its efficacy owing to potentially heightened binding affinity compared to that of the monomer, we synthesized the LTLRGLTLR peptide with an amidated C-terminus, designated as di-LTLR′. To assess the standalone effect of this dimeric peptide on bacterial growth, *P. aeruginosa* was exposed to various concentrations of di-LTLR′ in the absence of antibiotics, and cell viability was assessed. Importantly, di-LTLR′ exhibited bactericidal activity against *P. aeruginosa*, with an ED50 value of approximately 400 mM (Fig. 4), indicating a qualitative transformation of the peptide owing to dimerization. This unexpected discovery underscores the potential of this dimeric peptide as a novel antibiotic. However, considering the prospect of employing a new agent, it is imperative to investigate whether the dimeric peptide poses any adverse effects on mammalian cells.

**Fig. 4.**
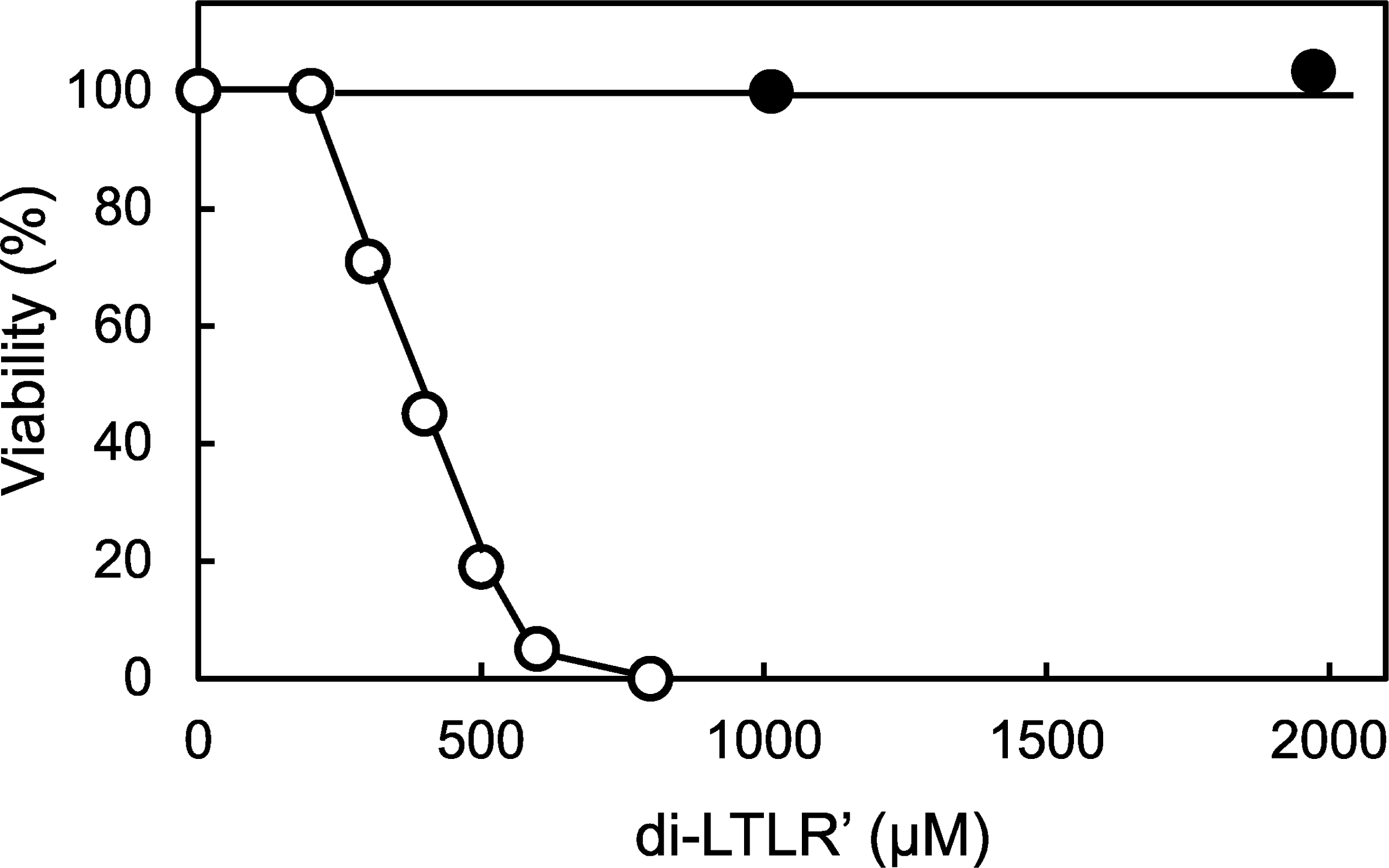
Activities of di-LTLR′ against *P. aeruginosa* and mammalian cells. *P. aeruginosa* PAO1 was incubated with di-LTLR′ at different concentrations in the absence of antibiotics at 37 °C for 4 h. Cell viability was determined using untreated cells as a control (○). Conversely, HeLa cells were cultured in medium containing 1 mM or 2 mM di-LTLR′, and cell viability was determined as described in the Materials and Methods section (●).

To address this concern, HeLa cells were cultured with the dimeric peptide, and cell viability was evaluated. As depicted in Fig. 4, di-LTLR′ (at concentrations of 1 or 2 mM) had no discernible impact on HeLa cell growth, suggesting a low risk of adverse side effects associated with this dimeric peptide.

### Activity of Dimeric Peptides Against MDR *P. aeruginosa*

We proceeded to evaluate the efficacy of the dimeric peptide against MDR *P. aeruginosa*. MDR *P. aeruginosa* cells were incubated with di-LTLR′, and subsequent cell viability assays were conducted. As depicted in Fig. 5, di-LTLR′ exhibited significant bactericidal activity against MDR *P. aeruginosa*, with an ED50 value of approximately 400 mM. Notably, the ED50 value of di-LTLR′ against the MDR strain mirrored that against the wild-type strain, indicating its capability to overcome multidrug resistance and effectively combat *P. aeruginosa*. This discovery holds considerable significance, given the pressing challenge posed by the emergence of MDR *P. aeruginosa* in clinical settings.

**Fig. 5.**
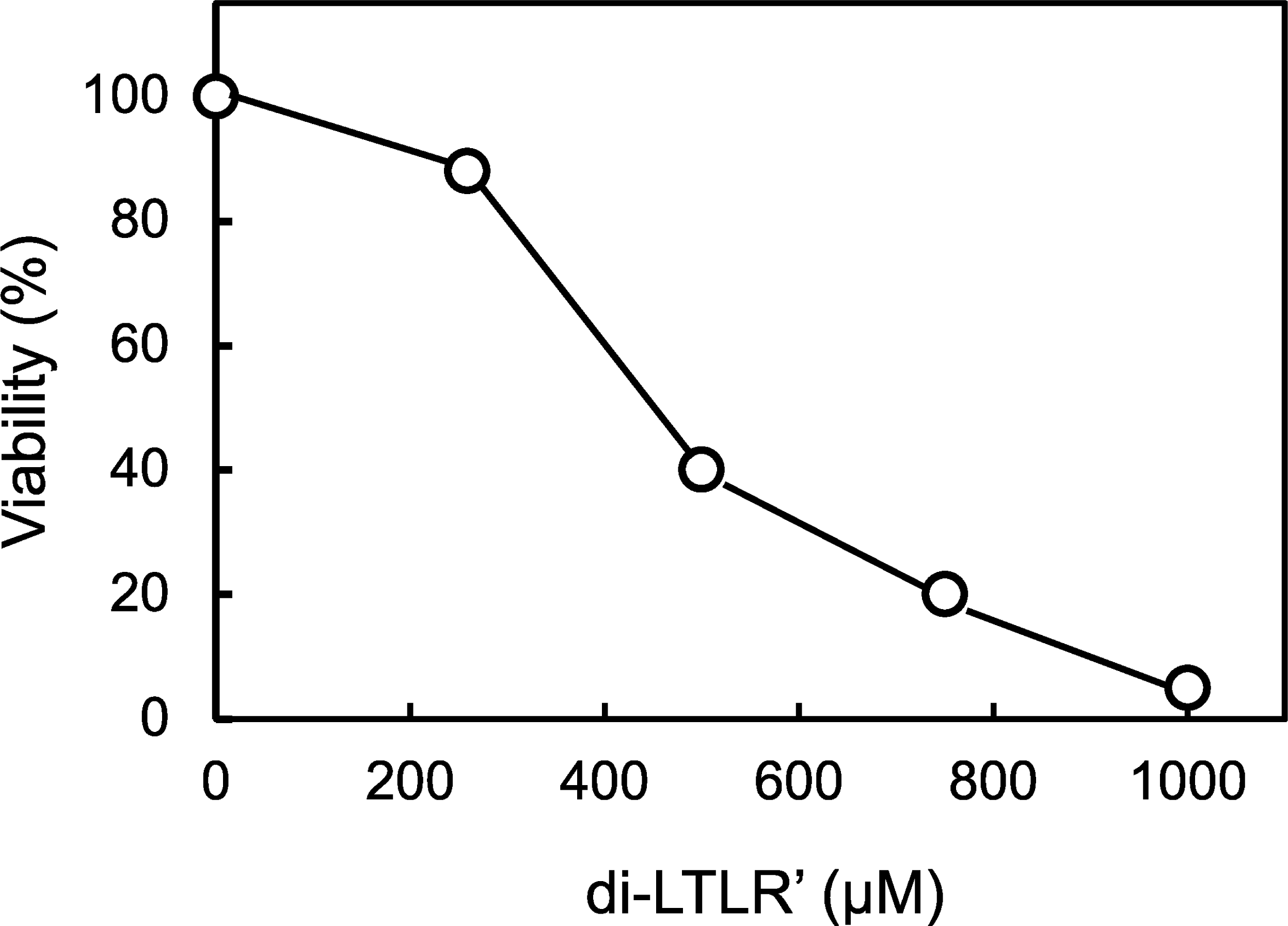
Bactericidal activity of di-LTLR′ against MDR *P. aeruginosa*. MDR *P. aeruginosa* strain was incubated with di-LTLR′ at different concentrations at 37 °C for 4 h. Cell viability was determined using untreated cells as controls.

### Peptide Activity Targeting the BAM Complex of *A. baumannii*

The demonstrated effectiveness of targeting the protein–protein interaction sites of the BAM complex in *P. aeruginosa* suggests a promising avenue for developing novel antibiotic agents, potentially extending to other Gram-negative bacteria. To investigate this premise, we focused on *A. baumannii*, a prominent MDR pathogen prevalent in hospital settings worldwide (18). Peptides designed to target the interface between BamA and BamB in the BAM complex of *A. baumannii* were employed. The identification of the amino acid sequence constituting the target site necessitated aligning BamB proteins from *Escherichia coli*, *P. aeruginosa*, and *A. baumannii* using ClustalW. As illustrated in Fig. 6, the amino acid sequence FSLR *in A. baumannii* BamB corresponded to LTLR in *P. aeruginosa* BamB. Subsequently, a dimeric peptide, FSLRGFSLR, with an amidated C-terminus (designated di-FSLR′), was synthesized.

**Fig. 6.**
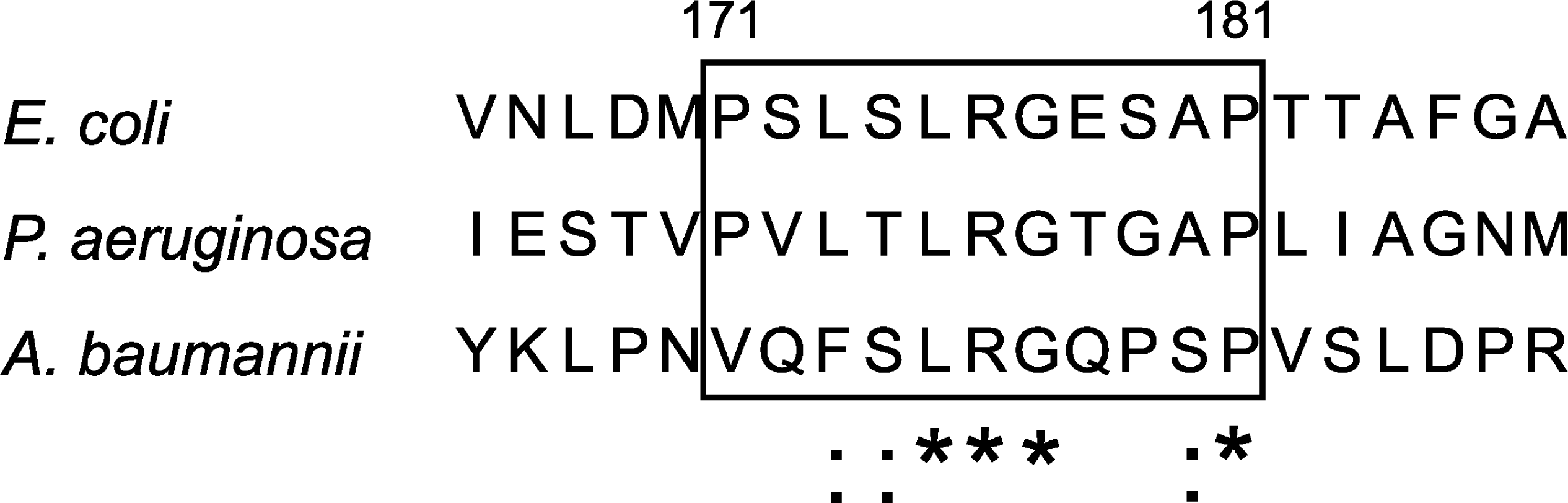
Homology of BamB regions interacting with BamA in *E. coli*, *P. aeruginosa*, and *A. baumannii*. Alignment of the three BamB proteins was performed using ClustalW, revealing homology between the region from residues 159 to 169 of *A. baumannii* BamB and the region from residues 171 to 181 of *E. coli* BamB, as well as the region from residues 159 to 169 of *P. aeruginosa* BamB.

Upon incubation with di-FSLR′ at varying concentrations, wild-type *A. baumannii* displayed dose-dependent bactericidal activity (Fig. 7), with an approximate ED50 value of 60 mM. This outcome underscores the efficacy of this strategy against *A. baumannii*. These findings strongly suggest the broad applicability of this approach across diverse Gram-negative bacterial strains.

**Fig. 7.**
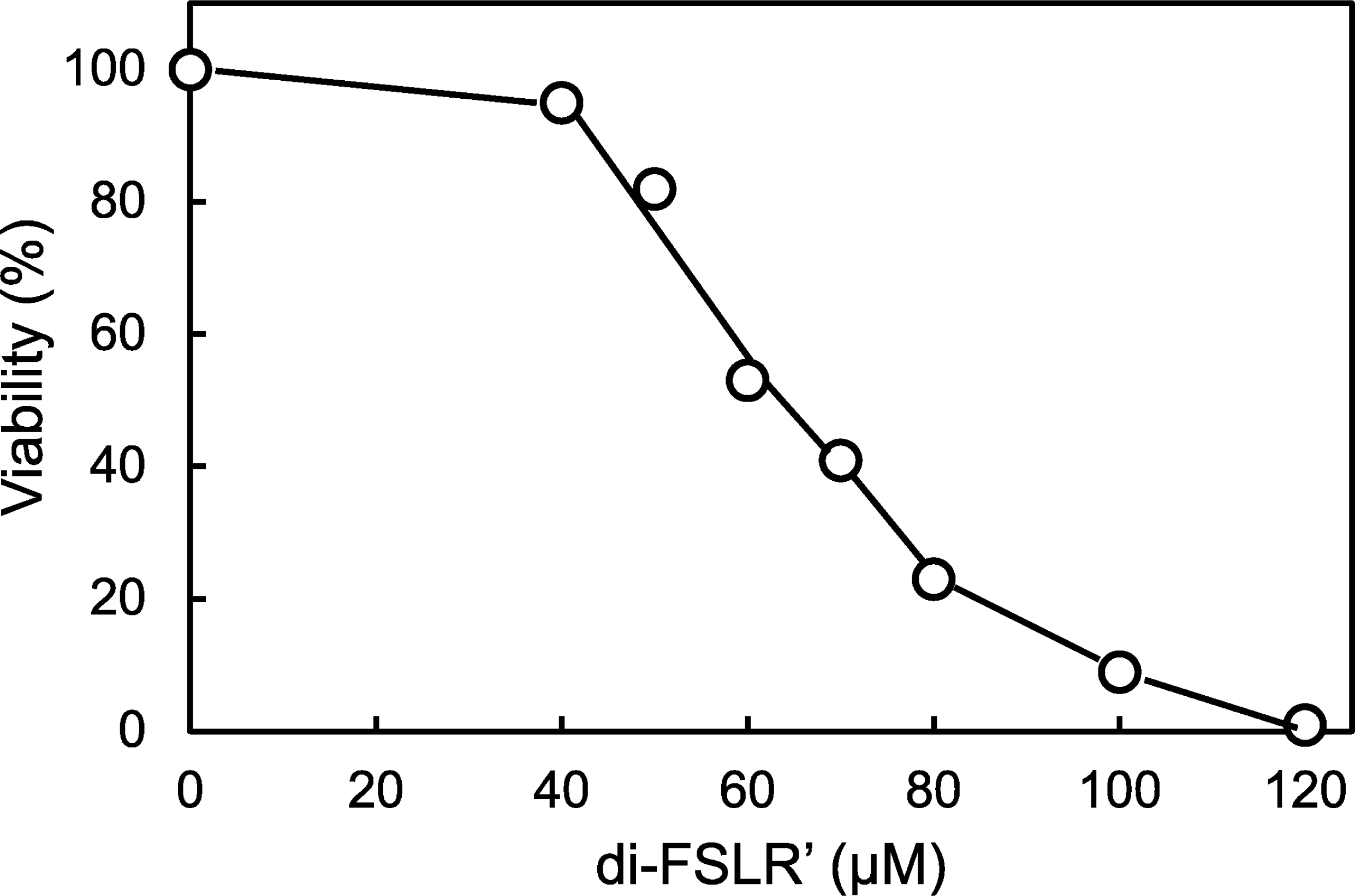
Bactericidal activity of di-FSLR′ against *A. baumannii*. Wild-type *A. baumannii* ATCC 19606 was incubated with di-FSLR′ at different concentrations and 37 °C for 4 h. Cell viability was determined using untreated cells as controls.

Notably, the ED50 value of di-FSLR′ against *A. baumannii* (approximately 60 mM) was considerably lower than that of di-LTLR′ against *P. aeruginosa* (approximately 400 mM), indicating a higher bactericidal activity of di-FSLR′ against *A. baumannii* than that of di-LTLR′ against *P. aeruginosa*.

### Activity of the Dimeric Peptide against MDR *A. baumannii*

To explore whether di-FSLR′ possesses bactericidal activity against MDR *A. baumannii*, MDR *A. baumannii* was incubated with di-FSLR′ at varying concentrations, and cell viability was assessed. As illustrated in Fig. 8, di-FSLR′ exhibited bactericidal activity against MDR *A. baumannii*, with an ED50 of approximately 50 mM. Importantly, the ED50 values of di-FSLR′ were comparable between the wild-type and MDR strain, indicating that this dimeric peptide can effectively combat bacterial resistance in *A. baumannii*. These findings suggest the potential utility of this peptide in the treatment of MDR *A. baumannii* infections.

**Fig. 8.**
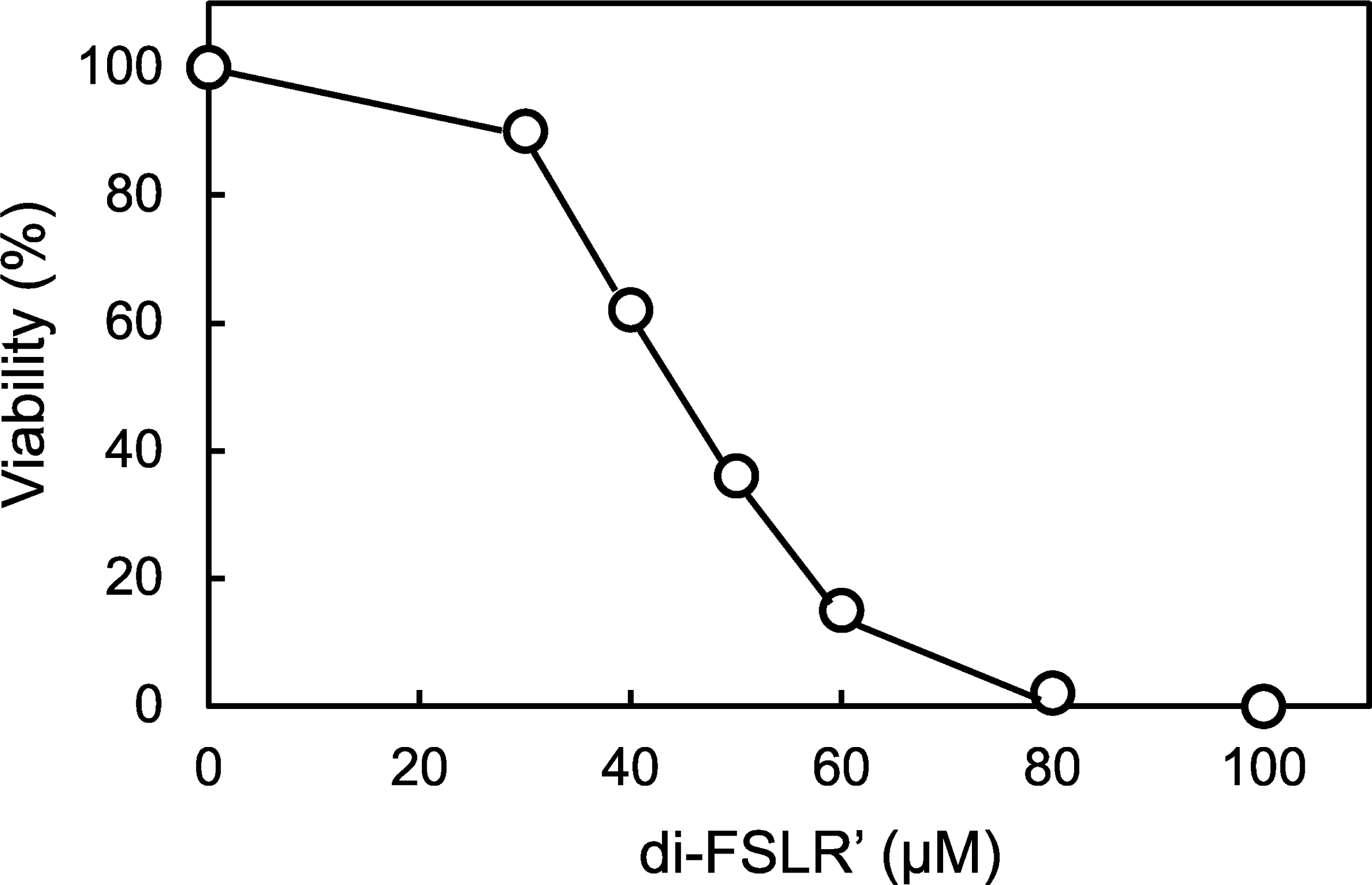
Bactericidal activity of di-FSLR′ against MDR *A. baumannii*. MDR *A. baumannii* strains were incubated with di-FSLR′ at different concentrations at 37 °C for 4 h. Cell viability was determined using untreated cells as controls.

### Activity of di-FSLR′ against *P. aeruginosa*

Our findings underscore the significance of disrupting interactions between Bam proteins for antimicrobial efficacy. Consequently, it is plausible that di-FSLR′ demonstrates bactericidal activity not only against *A. baumannii* but also against *P. aeruginosa*, given its homology to di-LTLR′. It is reasonable to assume that di-FSLR′, similar to di-LTLR′, could bind to the target site within the BAM complex of *P. aeruginosa*.

To validate this hypothesis, we treated *P. aeruginosa* PAO1 cells with varying concentrations of di-FSLR′ and assessed cell viability. As depicted in Fig. 9, di-FSLR′ exhibited bactericidal activity against *P. aeruginosa*, with an approximate ED50 value of 250 mM. Notably, this value was approximately four times higher than that observed for *A. baumannii*, indicating that while di-FSLR′ displayed bactericidal activity against *P. aeruginosa*, its potency was comparatively lower than that against *A. baumannii*. This observation aligns with expectations, as di-FSLR′ is expected to exhibit greater affinity for the target site of *A. baumannii* than that of *P. aeruginosa*.

**Fig. 9.**
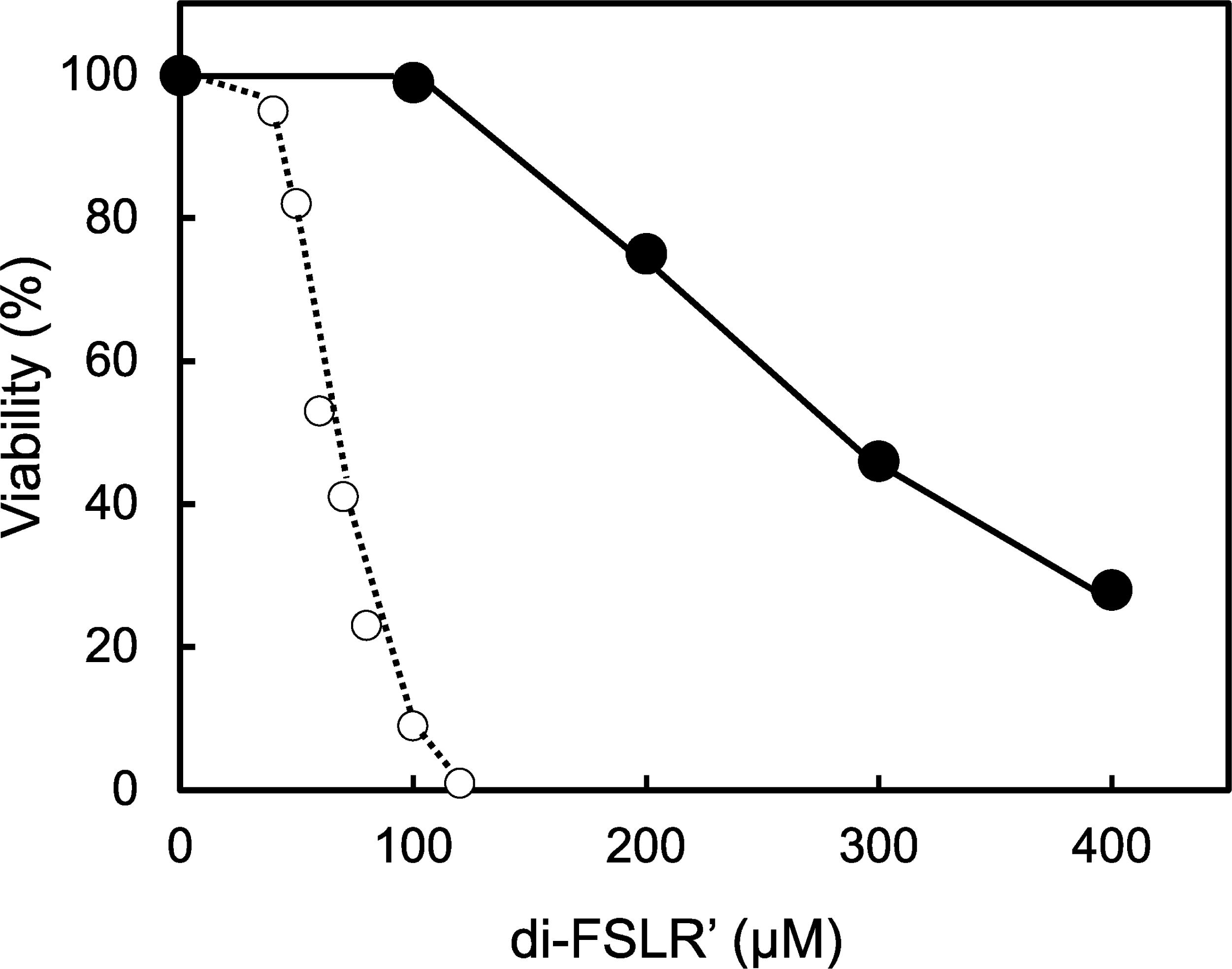
Bactericidal activity of di-FSLR′ against wild-type *P. aeruginosa*. *P. aeruginosa* PAO1 was incubated with di-FSLR′ at different concentrations at 37 °C for 4 h. Cell viability was determined using untreated cells as a control. The dotted line represents the bactericidal activity of di-FSLR′ against wild-type *A. baumannii*.

## DISCUSSION

A growing number of bacterial strains are developing resistance to multiple antibiotics currently in use; MDR Gram-negative bacteria are of notable concern in healthcare settings (1–4). Our aim was to develop novel antibiotic agents that are effective against MDR Gram-negative bacteria by targeting the BAM complex. Previously, we showed that the BAM complex could be a promising drug target by demonstrating that a peptide based on homologous sequences of BamD has adjuvant activity against *P. aeruginosa* (6). In this study, we targeted the interface between BamA and BamB in the BAM complex and demonstrated that these targeting peptides exhibit bactericidal activity against MDR *P. aeruginosa* and *A. baumannii*.

The LTLR peptide, one of the targeted peptides, exhibited no bactericidal activity against *P. aeruginosa*. However, it demonstrated antibiotic adjuvant properties, indicating its potential for the development of new antimicrobial agents.

We propose the following mechanism of action to explain the antibiotic adjuvant activity of the peptide. Upon binding to its target site, the peptide obstructs the interaction between BamA and BamB, resulting in impaired assembly and diminished function of the BAM complex. Consequently, the transport of outer membrane proteins is disrupted, compromising the integrity of the outer membrane, and reducing its barrier function, thereby facilitating the passage of antibiotics through the outer membrane. According to this proposed mechanism, the peptide is anticipated to act as an adjuvant irrespective of the antibiotic type used. Importantly, this peptide functions as an antibiotic adjuvant for PMB and OFLX.

Multidrug efflux pumps are pivotal in conferring multidrug resistance in Gram-negative bacteria. Given that the *P. aeruginosa* nalB strain exhibits high expression of the MexAB-OprM efflux pump, we investigated the efficacy of a specific peptide against *P. aeruginosa* nalB to assess whether this efflux pump can expel the peptide. Notably, our findings revealed that this peptide could serve as an adjuvant against *P. aeruginosa* nalB, indicating its non-interaction with multidrug efflux pumps.

During our investigation into the impact of peptide modification on activity, we discovered that the dimeric peptide di-LTLR′ demonstrated bactericidal activity against *P. aeruginosa*, contrary to our initial expectation. We originally hypothesized that peptide dimerization would enhance binding affinity to the target, thereby augmenting adjuvant activity. However, the disparity in activity between monomeric and dimeric peptides prompted further inquiry. We postulated that differences in blocking mechanisms between monomeric and dimeric peptides might underlie this variation in activity. Specifically, while monomeric peptides can bind to the target site, their blocking action may be insufficient to fully deactivate the BAM complex, resulting in compromised outer membrane integrity. By contrast, when the dimeric peptide binds to the target site, one unit of the dimer is engaged in target binding, while the other unit protrudes, causing steric hindrance in protein–protein interactions. Consequently, the function of the BAM complex is substantially inhibited, ultimately leading to cell death.

However, the discovery that the dimeric peptide demonstrates bactericidal activity, even against MDR strains of *P. aeruginosa*, holds considerable significance. Given the pressing need for novel agents to combat MDR *P. aeruginosa* infections, the dimeric peptide emerges as a potential treatment option.

Similarly, our investigation into di-FSLR′, a dimeric peptide designed to target the interface between BamA and BamB of *A. baumannii*, revealed its bactericidal activity against MDR *A. baumannii*. This finding is noteworthy owing to the widespread concern posed by MDR *A. baumannii* in healthcare settings worldwide.

The novel mode of action exhibited by these peptides suggests a lower likelihood of pre-existing resistance, supporting the assumption that they can exert bactericidal effects even against MDR strains of both *P. aeruginosa* and *A. baumannii*.

The emergence of drug-resistant bacteria is an inevitable consequence of developing and utilizing new antimicrobial agents for treating bacterial infections (19). Bacteria develop resistance to target peptides by introducing mutations into the target site to impede peptide binding. However, the occurrence of such mutations, which necessitates altering the interface structure while maintaining compatibility between BamA and BamB, is exceedingly rare. Furthermore, even if bacteria develop resistance to these peptides, this issue can be promptly addressed by designing targeting peptides adapted to mutated sites.

In this study, we showed that peptides targeting the interface between Bam proteins can possess bactericidal activity against MDR strains of *P. aeruginosa* and *A. baumannii*. These findings suggest that the targeting of such peptides may prove effective against other MDR Gram-negative bacteria. To develop more effective targeting peptides, the following peptides need to be created and changes in their antimicrobial activities need to be examined: (1) trimeric and tetrameric forms of the LTLR or FSLR peptide; (2) peptides containing sequences other than LTLR in the region of BamB interacting with BamA; and (3) cyclic targeting peptides. Furthermore, we will develop targeting peptides against other Gram-negative bacteria such as *Klebsiella pneumoniae*. The findings of our study pave the way for the development of novel antimicrobial agents against MDR Gram-negative bacteria.

## MATERIALS AND METHODS

### Bacterial Strains and Culture Conditions

The bacterial strains utilized in this study included *P. aeruginosa* PAO1 (standard strain), a *P. aeruginosa* nalB mutant overexpressing the MexAB-OprM efflux pump (provided by Dr. T. Nakae, Department of Molecular Life Science, Tokai University School of Medicine, Isehara, Kanagawa, Japan), the MDR *P. aeruginosa* 601 strain (provided by Dr. N. Koike, Department of Microbiology, Tokyo Medical University, Tokyo, Japan), *A. baumannii* ATCC 19606 (wild-type strain), and the MDR *A. baumannii* 10659 (provided by Dr. Y. Akeda, Research Institute for Microbial Diseases, Osaka University, Osaka, Japan).

Bacterial strains were cultured in Luria-Bertani (LB) broth composed of 10 g tryptone (Becton, Dickinson And Company, Sparks, MD, USA), 5 g yeast extract (Becton, Dickinson And Company), and 10 g NaCl per liter of water, adjusted to pH 7.0. Cultures were maintained at 37 °C and agitated at 150 rpm.

### Peptides

The peptides listed in Table 1 were synthesized and purified to >95% purity using high-performance liquid chromatography (Scrum Inc., Tokyo, Japan).

**TABLE 1.** List of synthesized peptides.

| Peptide | Amino acid sequence | Modification |
| --- | --- | --- |
| **LTLR** | LTLR | None |
| **LRTL** | LRTL | None |
| **LTLR'** | LTLR | C-terminal amidation |
| **di-LTLR'** | LTLRGLTLR | C-terminal amidation |
| **di-FSLR'** | FSLRGFSLR | C-terminal amidation |

### Antibiotics

PMB and OFLX were procured from Sigma–Aldrich (St. Louis, MO, USA) and LKT Laboratories (St. Paul, MN, USA), respectively.

### Measurement of Antimicrobial Activity

Bacterial overnight cultures were diluted to 1:500 in LB broth and incubated at 37 °C. Bacteria in mid-log phase growth were exposed to peptides at various concentrations (0 – 3 mM) or left untreated for 4 h at 37 °C. Following treatment, bacteria were diluted with PBS by more than 1000 times and plated onto LB agar plates. After overnight incubation at 37 °C, bacterial colonies were counted. Cell viability was calculated as follows: (number of colonies from treated bacteria/number of colonies from untreated bacteria) × 100. Data represent average values from at least two independent experiments.

### Measurement of Cytotoxicity

Cytotoxicity testing of the peptides was conducted as follows: HeLa cells (approximately 10,000 cells) were seeded in 96-well plates containing 200 µl of DMEM/Ham’s F-10 medium supplemented with 10% fetal bovine serum, 0.006% penicillin, and 0.01% streptomycin and then cultured at 37 °C for 4 h under a humidified atmosphere with 5% CO2. After peptide or phosphate buffer addition, the cells were further incubated for 72 h under the same conditions.

Cytotoxicity was measured using Alamar Blue reagents (Bio-Rad Laboratories, Hercules, CA, USA) by spectrophotometry. Cell viability was determined by comparing the absorbance of treated cells with that of control cells.

## ACKNOWLEDGMENTS

We are grateful to Dr. T. Nakae, Dr. N. Koike, and Dr. Y. Akeda for providing the bacterial strains.

## AUTHOR CONTRIBUTIONS

**Eisaku Yoshihara**: Conceptualization, Investigation, Methodology, Writing-original draft, Writing-review and editing. **Hidefumi Kakizoe**: Investigation. **Kazuo Umezawa**: Writing-review and editing. **Kentaro Wakamatsu**: Writing-review and editing. **Satomi Asai**: Conceptualization, Supervision, Writing-review and editing.

## DATA AVAILABILITY

Not Applicable.

